# ALDH2 Deficiency Drives Proliferative Mammary Morphogenesis and Epithelial Cell Stemness via Oxidative Stress and Estrogen Receptor Activation

**DOI:** 10.1101/2025.09.25.678642

**Authors:** Zhikun Ma, Amanda B Parris, Miles Lester, De’ja Gissendanner, Vasilis Vasiliou, Xiaohe Yang

## Abstract

Alcohol consumption has been linked to breast cancer, partly due to the accumulation of toxic aldehydes like acetaldehyde, a carcinogenic byproduct of ethanol metabolism. ALDH2, a key mitochondrial enzyme, detoxifies acetaldehyde and other harmful aldehydes that drive oxidative stress, DNA damage, and hormonal dysregulation, key factors in carcinogenesis. Despite the possible link between alcohol consumption and breast cancer risk, little is known about how ALDH2 deficiency itself, independent of alcohol exposure, affects mammary gland biology and cancer susceptibility. Genetic variants leading to ALDH2 deficiency are highly prevalent in East Asian populations, where individuals carrying inactive ALDH2 alleles experience impaired aldehyde detoxification. While these individuals are at increased risk for alcohol-related cancers, the impact of ALDH2 deficiency on mammary gland development and homeostasis in the absence of alcohol exposure remains unexplored. To address this, we utilized a C57BL/6-based ALDH2 knockout (*Aldh2-/-*) mouse model to investigate its effects on mammary proliferation and development. Our findings revealed that *Aldh2-/-* mice exhibited hyperproliferative mammary glands, characterized by increased epithelial cell density, ductal expansion, and elevated Ki67+ cells. Flow cytometry analysis demonstrated a rise in luminal and basal epithelial subpopulations, alongside enhanced mammary epithelial stemness, as evidenced by increased mammosphere formation and colony-forming efficiency. At the molecular level, ALDH2 deficiency activated oxidative stress pathways, marked by elevated 8-OHdG, p38 MAPK, NF-κB, and Nrf2 signaling, alongside DNA damage responses involving p53 and H2A.X. Importantly, we also identified a previously unrecognized upregulation of RANKL in Aldh2-/- mammary glands, implicating the RANK/RANKL axis as a critical mediator linking aldehyde stress to NF-κB/p38 MAPK activation and enhanced mammary stemness. Furthermore, hormonal dysregulation was observed, with a significant increase in ERα and PR expression and phosphorylation. Dysregulated ER signaling was further linked to enhanced erbB3 activation and downstream signaling, including the cyclin D1-pRb-E2F1 axis. Our results suggest that the accumulation of endogenous aldehydes, independent of alcohol exposure, profoundly alters mammary morphogenesis, epithelial repopulation, and stemness. Mechanistically, this occurs through oxidative stress activation and DNA damage pathways, leading to metabolic changes and upregulation of estrogen receptor and receptor tyrosine kinase signaling. This study highlights for the first time the potential role of ALDH2 deficiency in increasing mammary tissue susceptibility to oncogenic factors and breast cancer risk.

## 1. Introduction

Aldehyde dehydrogenase 2 (ALDH2) plays a critical role in the detoxification of aldehydes, particularly acetaldehyde, and is essential for maintaining cellular homeostasis [1]. Importantly, this enzyme is not only involved in the metabolic pathways arising from alcohol consumption and environmental toxins, it is also essential for the cleaning of endogenous aldehydes generated through the metabolism of amino acids, alcohol-related substances, and lipid peroxidation [2, 3]. It is critical to understand the impact of ALDH2 function relative to endogenous aldehydes in different tissues.

The *ALDH2* polymorphism, particularly the *ALDH2*2* (rs671) allele, is associated with a dramatic reduction in ALDH2 enzyme activity [4]. This polymorphism is highly prevalent in East Asian populations, with an estimated allele frequency of 30-50% [5]. Epidemiological studies indicate that *ALDH2* polymorphisms significantly influence susceptibility to various diseases, including cancers [1, 6]. Increasing evidence suggests that the functional deficiency associated with *ALDH2* polymorphisms and the resulting elevated aldehyde levels may promote the development and progression of esophageal, head and neck, liver, and colorectal cancers [7-10]. The relationship between *ALDH2* polymorphisms and breast cancer risk remains complex. While some studies support a correlation between *ALDH2* polymorphisms and an increased risk of breast cancer, particularly in specific populations [11, 12], other research finds no significant association [13, 14]. This underscores the need for further investigation to draw definitive conclusions. Studies using preclinical models with well-controlled experimental conditions are especially needed to advance understanding of these critical issues. However, reports in this area are limited.

Extensive biochemical studies indicate that ALDH2 is important for mitigating oxidative stress associated with cellular metabolism [15, 16]. As a mitochondrial enzyme, ALDH2 aids in the detoxification of harmful aldehydes produced during metabolic processes [17]. Deficiency in ALDH2 can increase oxidative stress in cells, which is linked to various pathological conditions, including cardiovascular diseases and complications related to alcohol metabolism [18-21]. Fully elucidating the diverse roles of ALDH2 in cellular metabolism and its interaction with oxidative stress pathways with molecular insight is critical for understanding its potential interventions for oxidative stress-related diseases. However, most investigations focus on the liver, cardiovascular and other tissues [16, 18, 22, 23], with limited evidence regarding the impact of ALDH2 deficiency on mammary gland physiology. Existing literature highlights the utility of *Aldh2-/-* mouse models in elucidating the *in vivo* roles of ALDH2 across different organs, such as liver and brain [22, 24-26]. However, *in vivo* studies on ALDH2’s function on mammary development and the underlying mechanisms have not been explored.

The biochemical functions of ALDH2 and its role in mitochondrial function, oxidative stress, and DNA damage have been extensively studied [15, 16]. However, most investigations focus on the liver and other tissues [16], with limited evidence regarding the impact of ALDH2 deficiency on mammary gland physiology. Existing literature highlights the utility of *Aldh2-/-* mouse models in elucidating the *in vivo* roles of ALDH2 across different organs, such as liver and brain [22, 24-26]. However, the implications of ALDH2’s function on mammary development have not been explored.

Mammary gland development is susceptible to various environmental and endogenous influences that can alter cellular dynamics, particularly with respect to breast cancer risk [27]. Previous findings indicate that aberrations in mammary epithelial cell proliferation and differentiation often precede the onset of breast cancer [28]. Recent advances suggest that mammary stem cells, which are pivotal for mammary tissue development and differentiation, are particularly vulnerable to metabolic disruptions [29, 30]. Based on the cancer stem cell theory, mutated or dysregulated mammary stem/progenitor cells are likely the origin of cancer stem cells [31]. Changes in stem/progenitor cell dynamics, which are reflected in mammary epithelial repopulation and functional alterations in their stemness, are emerging biomarkers for the evaluation of mammary proliferation and differentiation in premalignant tissues [32]. Considering the biochemical changes associated with ALDH2 deficiency observed *in vitro* [15, 16], it is imperative to examine its *in vivo* effects on mammary development and its potential implications for breast cancer risk.

In this study, we utilized the C57BL/6-*Aldh2*-/- mouse model to investigate the consequences of ALDH2 deficiency on mammary development. Our findings reveal that ALDH2 knockout (KO) induces considerable proliferative dynamics, morphological changes and modified signaling pathways, within mammary epithelial cells, implicating alterations that may heighten breast cancer risk under various conditions. Central to our observations were robust changes in the repopulation dynamics of mammary epithelial cells, suggesting a critical role for ALDH2 in regulating the stem cell characteristics of these cells, which will advance our understanding of the potential pathways through which ALDH2 may influence breast cancer etiology.

## 2. Materials and Methods

### 2.1 Reagents

p-p38 MAPK (Thr180/Tyr182) (4511), p38 MAPK (9212), p-NF-κb (Ser536) (3033), p-NF-κb (8242), Nrf2 (12721), p-H2A.X (Ser139) (9718), H2A.X (2595), PR (8757), p-PR (Ser190) (3171), p-EGFR (Ser1046/1047) (2238), p-erbB-3 (Tyr1197) (4561), p-Akt (Ser473) (4060), p-Erk1/2 (Thr202/Tyr204) (9101), p-ERα (Ser118) (2511), c-Myc (5605), p-Rb (Ser807/811) (8516), Rb (9309) and GAPDH (5174) primary antibodies were sourced from Cell Signaling Technologies. Primary antibodies against ALDH2 (sc-100496), p53 (sc-126), MDM2 (sc-965), Bcl-2 (sc-7382), cyclin D1 (sc-246), Akt1 (sc-5298), Erk2 (sc-1647), ERα (sc-8002), ERβ (sc-8974), EGFR (sc-373746), erbB-3 (sc-285), E2F1 (sc-251) and β-actin (sc-47778) were purchased from Santa Cruz Biotechnology. Anti-rabbit (7074) and anti-mouse (7076) HRP-linked secondary antibodies were from Cell Signaling Technologies. For immunohistochemistry, the Ki67 (PA5-19462) antibody was purchased from Invitrogen, 8-OHdG (sc-66036) was purchased from Santa Cruz Biotechnology and the secondary anti-rabbit antibody for immunohistochemistry was part of the Vecta-Stain ABC kit (Vector Labs). Flow cytometry antibodies against CD16/32 (553141), CD49f (555735), CD24 (553260), and CD61 (553345) were purchased from BD Biosciences, and those against CD31 (102508), CD45 (103106), and Ter-119 (116208) were purchased from BioLegend.

### 2.2 Animals and husbandry

Female C57BL/6 mice were purchased from Jackson Laboratory. The C57BL/6-*Aldh2-/-* mice were provided by Dr. Vasilis Vasiliou’s laboratory at Yale University. The mice were maintained on base diet and housed in a temperature-controlled room with a 12-hour light-dark cycle. A total of 15 mice were allocated to each group for various analyses, including wholemount/histopathology, Western blotting, RNA extraction, mammosphere and colony-forming cell (CFC) assays, and flow cytometry, as detailed in the respective assay protocols. All procedures involving mice were performed with the approval of the Institutional Animal Care and Use Committee.

### 2.3 Whole Mount Analysis

The #4 inguinal mammary glands were dissected from control and *Aldh2-/-* mice at the endpoints and spread on slides. After fixation overnight in Carnoy’s solution, the glands were rehydrated through ethanol washes. Subsequently, they were stained with carmine alum, dehydrated with ethanol, cleared using xylene, and finally mounted with Permount, following established protocols [33]. The wholemount images were captured using a Nikon microscope. The complexity of the mammary ductal trees was assessed by quantifying the number of lateral buds in an area of 10 mm^2^, which was statistically analyzed using a Student’s *t*-test based on 5 mice from each group.

### 2.4 Histology and Immunohistochemistry

The harvested mammary tissues were fixed in formalin and processed for routine paraffin embedding/tissue sectioning. For H&E sections, the tissues were stained with hematoxylin and eosin. For immunohistochemistry (IHC), the VECTASTAIN Elite ABC kit from Vector Laboratories was used. Briefly, the slides were incubated with primary antibodies with appropriate dilutions (Ki67 1:5000, 8-OHdG 1:300, p-ERα 1:100 and cyclin D1 1:100) at 4°C overnight, followed by incubation with a biotinylated secondary anti-rabbit antibody and ABC reagent. Diaminobenzidine (DAB) was used for color development, and the slides were counterstained with hematoxylin [33]. Imaging and documentation were acquired using a Nikon Microscope.

### 2.5 MEC Isolation and Flow Cytometry

The #4 inguinal mammary glands from 16-week-old mice were collected and homogenized with a tissue chopper. The processed tissues were then digested with collagenase and hyaluronidase for 2 h at 37^°^C, as in our previous reports [34]. The resulting organoids were further digested with trypsin and dispase/DNase I and then filtered through a 40-μm mesh strainer. The single cell suspension of MECs was then used for flow cytometry and functional assays described below. For flow cytometry analysis, 1 x 10^6^ primary mammary epithelial cells underwent staining with fluorescent antibodies targeting lineage and cell surface markers, as previously described [34]. In brief, cells were first stained with anti-CD16/CD32 to block Fc receptors. PE-conjugated anti-CD31, anti-TER-119, and anti-CD45 were used to exclude endothelial, erythroid, and leukocyte cells, respectively. Additionally, biotin–streptavidin–APC anti-CD24, biotin–streptavidin–APC anti-CD61, and FITC-conjugated anti-CD49f were employed. The dead cells were excluded for subpopulation analysis by gating with 7-AAD positive cells. Lin^-^ CD24/CD49f cells were gated for stromal, luminal and basal subpopulations. Lin^-^ CD61/CD49f was used for the identification of progenitor enriched cells, as indicated by P2 subpopulation in Fig. 2C. FlowJo analysis software was used for gating and quantification of the individual cell populations.

### 2.6 Colony-forming cell assay

Single cell suspensions of the primary mammary epithelial cells were plated at a density of 4 × 10^3^ cells per 60-mm plate and incubated in a 5% CO_2_ atmosphere. Following a 10-day incubation period, colonies were rinsed with PBS, fixed in acetone and methanol (1:1), and stained using Wright’s Giemsa [34]. Imaging was performed using the Nikon SMZ 745T microscope and Nikon Elements Imaging System software.

### 2.7 Mammosphere assay

Isolated primary mammary epithelial cells were seeded at a density of 2.5 × 10^4^ cells per well in ultra-low attachment 24-well plates. The cells were cultured in the EpiCult-B Mouse Media (Stemcell Technologies) supplemented with 10 μg/ml insulin (Sigma), 1 μg/ml hydrocortisone (Sigma), 1× B-27 (Thermo Scientific), 20 ng/ml EGF (Stemcell Technologies), 20 ng/ml basic fibroblast growth factor (Stemcell Technologies), 4 μg/ml heparin (Stemcell Technologies), and 50 μg/ml gentamycin for 7 days. The primary mammospheres > 30 μm were counted and imaged. For secondary sphere formation assays, the cells of the primary spheres were trypsinized, and the resulting single-cell suspension was replated at a density of 1 × 10^3^ cells per well under identical conditions to form secondary spheres. After an additional 7 days of incubation, secondary mammospheres were counted and imaged for analysis [34]. The data from primary and secondary mammosphere assays in triplicate were analyzed with Student’s *t*-test.

For the assessment of anchorage-independent cell growth with 3D culture assay, primary mammary epithelial cells were seeded at a density of 1.5 × 10^4^ cells per well in matrigel in 48-well plates and cultured for 10 days. Subsequently, the colonies formed in the semi-solid media were stained with crystal violet, quantified and imaged. The data sets in triplicate were statistically analyzed with Student’s *t*-test.

### 2.8 Western blotting

The collected mammary tissues were homogenized in lysis buffer as previously reported [34]. Protein concentrations of the lysates were determined using a BCA Protein Assay kit (Thermo Scientific). Fifty (50) μg of protein lysate from each sample was separated using SDS-PAGE. The proteins on the gels were transferred to nitrocellulose membranes and blocked with 5% milk in Tris-buffered saline with Tween 20 (TBST). The membranes were then incubated overnight at 4°C with primary antibodies diluted in 5% bovine serum albumin in TBST. After washing with TBST, the membranes were incubated with HRP-labeled secondary antibodies at room temperature for 1 hour, followed by detection with enhanced chemiluminescence reagents (Thermo Scientific) [33]. Images of the protein bands were captured using an Azure imaging system.

### 2.9 RNA isolation and quantitative real-time PCR

Mammary tissues from indicated groups were harvested and snap-frozen in liquid nitrogen. Subsequently, the tissues were homogenized for RNA extraction utilizing the RNeasy Mini Kit (Qiagen). Total RNA (1 μg) from each sample was reverse transcribed with the iScript cDNA Synthesis Kit (Bio-Rad). Quantitative real-time PCR was performed with the Bio-Rad CFX 96TM Real-Time PCR System (Bio-Rad). The primers used in this study are summarized in Supplementary Table S1. Relative mRNA levels were quantified compared to control samples, utilizing cycle threshold (Ct) values and normalization to the GAPDH internal control as previously reported [34]. The differences among the compared groups were analyzed with Student’s *t*-test.

## 3. Results

### 3.1 ALDH2 KO induces proliferative mammary glands

The effect of ALDH2 deficiency on mammary development has not been documented. To investigate the impact of ALDH2 KO on mammary gland development and morphogenesis, we performed histopathological characterization of the mammary tissues from control and *Aldh2-/-* mice. Our preliminary data indicated that the changes at 16 weeks of age represented typical alterations associated with different *Aldh2-/-* status. Wholemount analysis showed that mammary tissues in the *Aldh2-/-* mice at this age exhibited more complex ductal growth (Fig. 1A), as indicated by increased lateral branching as compared to the control (Fig. 1B). H&E staining of the mammary tissues from *Aldh2-/-* mice indicated increased ductal thickness and proliferative morphology as compared to control (Fig. 1C). Morphogenic analysis indicated that ALDH2 KO induces proliferative mammary glands. This was further confirmed by Ki67 staining of the mammary tissues (Fig. 1D). The percentage of Ki67 positive cells was significantly higher in the *Aldh2-/-* mice than the control (Fig. 1E). Taken together, our results demonstrate that ALDH2 KO induces mammary epithelial proliferation and drives morphogenic changes in the process of glandular development.

**Figure 1.**
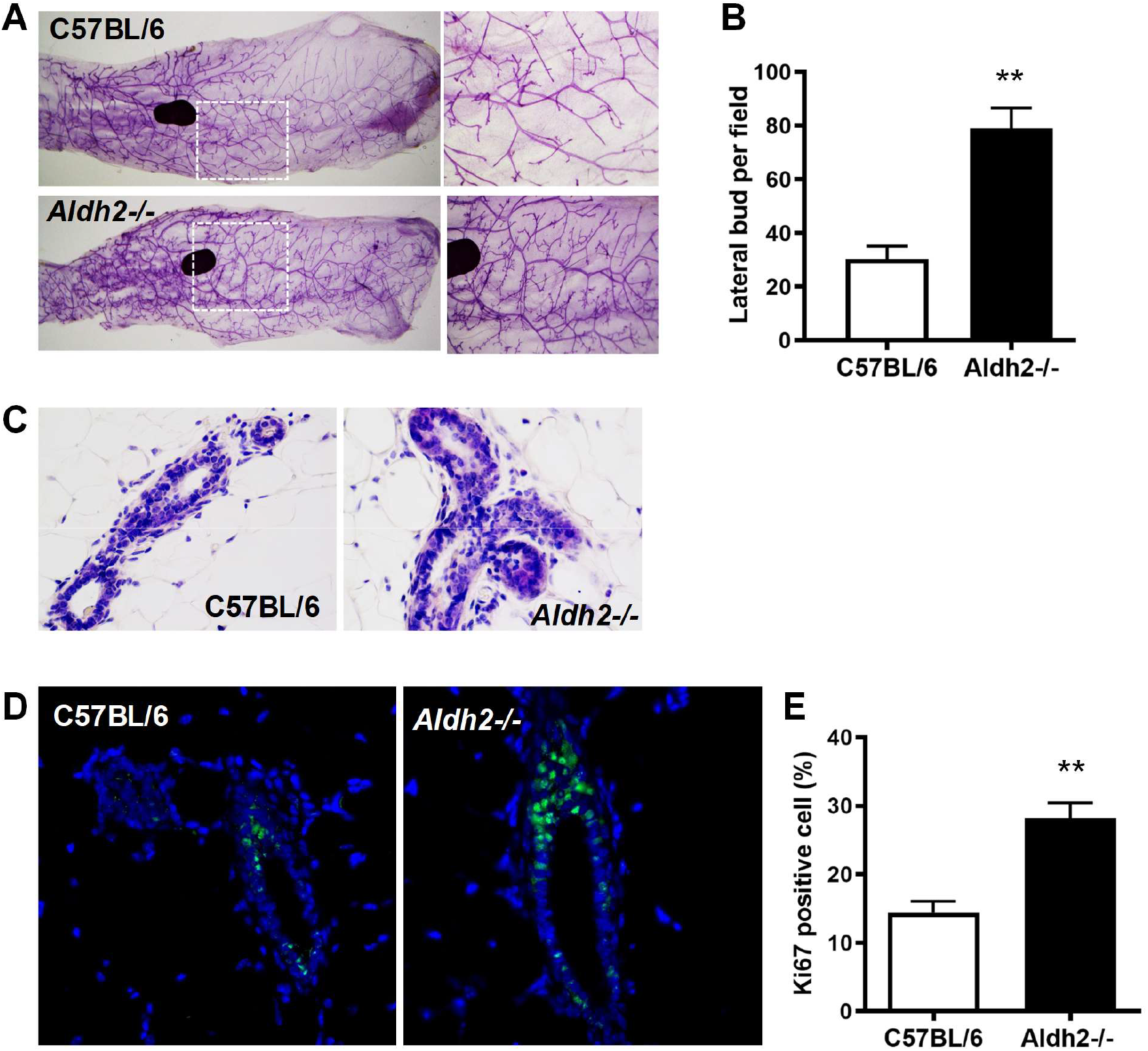
ALDH2 KO induces proliferative mammary glands of C57BL/6 mice. Mammary glands were collected for wholemount and histopathology analysis at 16 weeks of age. **A)** Representative mammary wholemounts of control and *Aldh2-/-* mice. Lateral bud numbers per 10 mm^2^, based on 5 mice of each group, were quantified in (B). **C)** H&E staining of mammary tissues with ductal structures. **D & E**) Ki67 staining of mammary tissues of control and *Aldh2-/-* mice. Ki67+ cells in green were detected with immunofluorescence (D). The percentage of Ki67+ mammary epithelial cells was quantified in E. (** p < 0.01).

**Figure 2.**
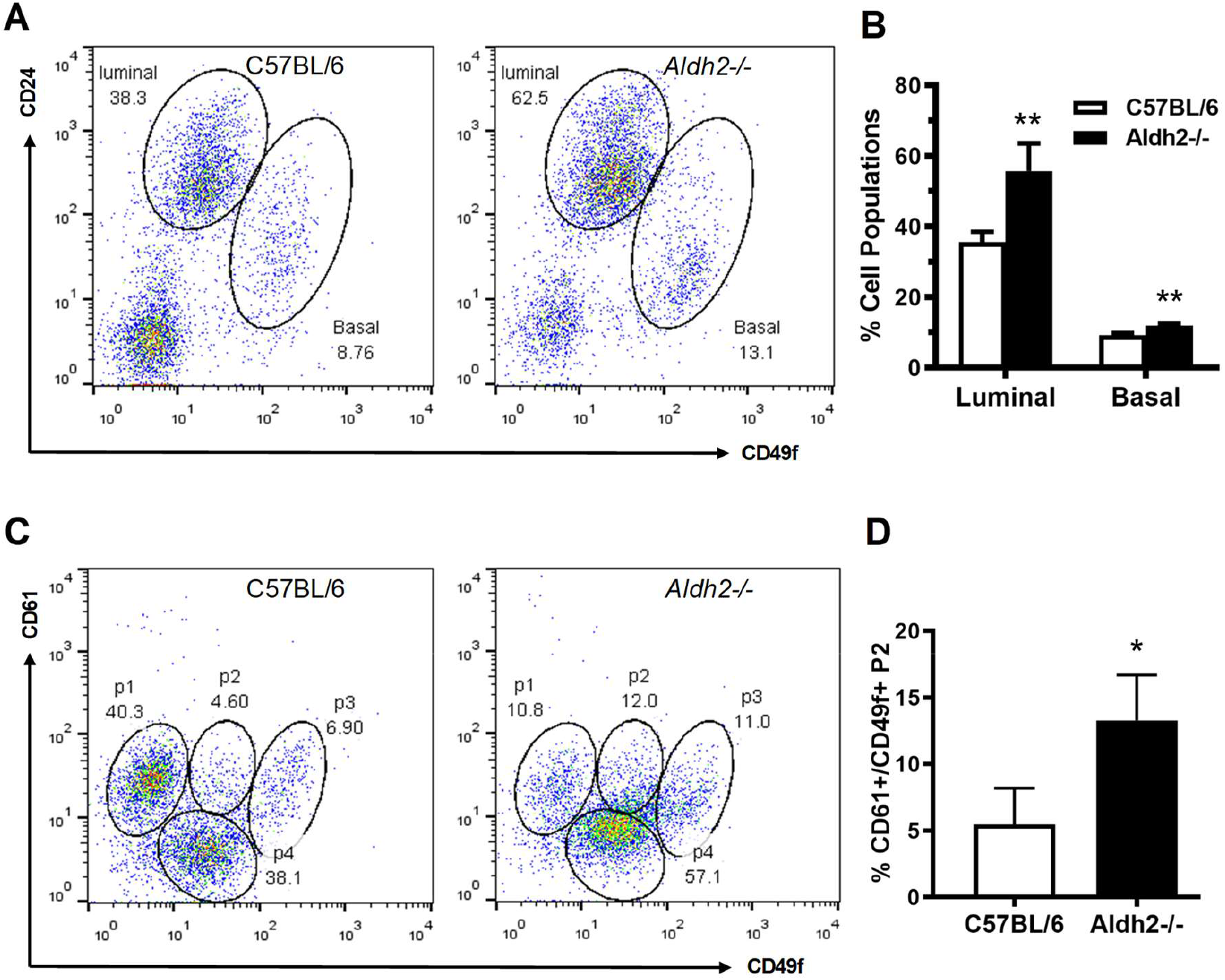
ALDH2 KO induces expansion of both luminal and basal mammary epithelial cell subpopulations with remarkable increase in luminal progenitor cells. Mammary cells isolated from control and *Aldh2-/-* mice were analyzed for different mammary epithelial cell subpopulations with flow cytometry. CD24 and CD49f were used to detect the relative composition of luminal, basal and stromal subpopulations (A-B). CD61 and CD49f were used to detect luminal progenitor cells (CD61+/CD49f+) enriched subpopulation (C-D). **A)** Representative plots of flow cytometry analysis based on CD24 and CD49f for luminal and basal subpopulations. **B)** Percentages of luminal/basal subpopulations from four mice of each group were statistically analyzed; **C)** Representative plots from the detection of subpopulation 2 (P2) of CD61+/CD49f+ cells in mammary tissues of control and *Aldh2-/-* mice. **D)** Quantification of CD61+/CD49f+ P2 cells of each group as in B. (** p < 0.01, *p < 0.05).

### 3.2 ALDH2 KO induces expansion of both luminal and basal mammary epithelial cell subpopulations and increase in luminal progenitor cells

Recent advances enable us to assess mammary epithelial cell proliferation and stemness with the characterization of mammary cell subpopulations using flow cytometry [35]. To understand the cellular mechanism underlying ALDH2 KO-induced proliferative mammary tissues, we examined the relative composition of luminal, basal and stromal subpopulations of mammary epithelial cells with CD24 and CD49f-based analysis. The results showed that the percentages of both basal and luminal cells, in particular of the *Aldh2-/-* mice, were significantly increased as compared to the control, suggesting that ALDH2 deficiency promotes the mammary gland’s developmental and/or physiological state (Fig 2A & B).

Moreover, CD61 is a recognized marker for luminal progenitor cells, commonly used in previous studies [36, 37]. In our CD61/CD49f-based analysis of mammary epithelial cells, the P2 subpopulation indicates an enrichment of luminal progenitor cells (Fig. 2C), which was significantly increased in the mammary tissues of *Aldh2-/-* mice. Our results indicate that the relative composition of luminal progenitor cells was notably higher in *Aldh2-/-* mice compared to controls (Fig. 2D). Taken together, the data from flow cytometry analysis suggests that ALDH2 KO promotes the expansion of both luminal and basal subpopulations, and the enrichment of luminal progenitor cells, which reflects alterations in cellular hierarchy associated with the proliferative state of mammary glands in *Aldh2-/-* mice.

### 3.3 ALDH2 KO promotes mammary epithelial cell stemness

The ALDH2 KO-associated expansion of basal and luminal subpopulations, along with progenitor-like cells, suggests that ALDH2 deficiency induces dynamic changes in mammary tissue stemness. We therefore examined its functional impact on mammary epithelial cell stemness through colony-forming cell (CFC) assays, mammosphere formation, and 3D culture. As shown in Fig. 3A, the number of CFCs, a functional readout of luminal and basal progenitor cell enrichment [38], was significantly increased in the mammary tissues of *Aldh2-/-* mice. Mammosphere formation assays, which detect the self-renewal potential of mammary stem cells [38], indicated that the mammosphere formation efficiency of cells derived from *Aldh2-/-* mice, including both primary and secondary spheres, was significantly higher compared to the control (Fig. 3B). Additionally, we performed semi-solid 3D culture to assess the anchorage-independent growth potential of mammary cells from both groups. As shown in Fig. 3C, the number of colonies in the *Aldh2-/-* group was significantly greater than in the control. Together, these results suggest that ALDH2 deficiency promotes mammary stemness, supporting our earlier observations of proliferative morphogenesis and the expansion of basal and luminal subpopulations of mammary epithelial cells.

**Figure 3.**
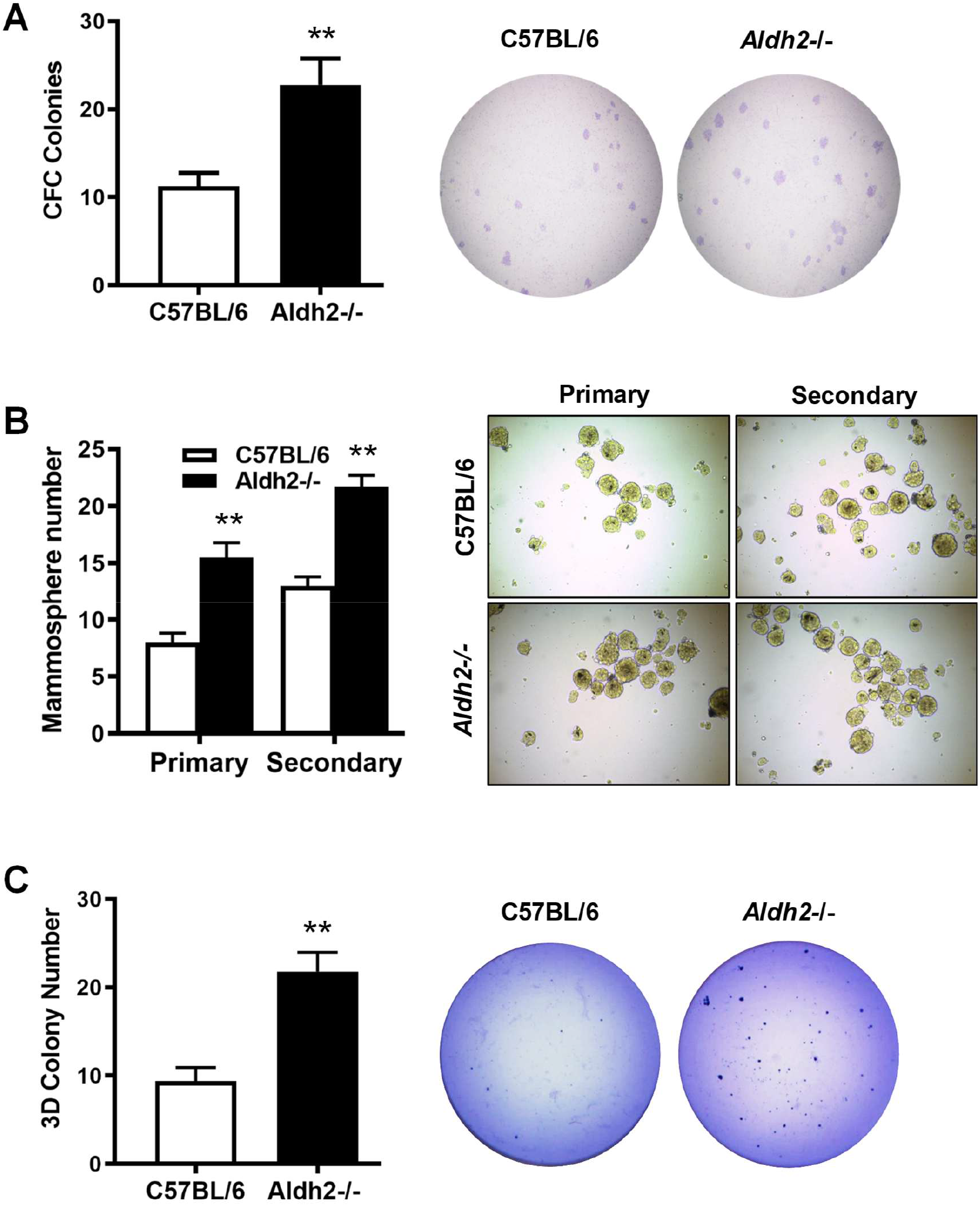
ALDH2 KO promotes mammary epithelial cell stemness. Primary mammary epithelial cells were isolated from control and *Aldh2-/-* mice (5 mice/group) and evaluated with functional assays for epithelial cell stemness. **A)** Detection of relative colony-forming cell (CFC) numbers in the mammary tissues of control and *Aldh2-/-* mice with CFC assays. CFC colonies formed in a dish inoculated with 4×10^3^ mammary cells from each group were documented and quantified. **B)** Mammosphere formation efficiency of mammary epithelial cells from control and *Aldh2-/-* mice. Primary mammary epithelial cells were cultured in supplemented-Epicult-B media for 7 days for the evaluation of primary mammospheres. Single cells from the primary spheres were cultured under the same conditions for secondary mammospheres. Spheres > 30 μm were counted and quantified for statistical analysis. **C)** Anchorage-independent growth of MECs was assessed with a 3D culture assay. Primary MECs from each group were grown in matrigel for 10 days, followed by fixation, staining, and colony analysis. Each of the assays was performed in triplicate (** p < 0.01).

### 3.4 ALDH2 KO activates oxidative stress signaling involving p38 MAPK and NF-κB and induces DNA damage pathway in mammary tissues

Previous studies have shown that ALDH2 KO induces the accumulation of toxic aldehydes, resulting in increased oxidative stress that activates various oxidative stress pathways in other tissues [21]. To understand the molecular mechanisms underlying altered mammary development in *Aldh2-/-* mice, we examined oxidative stress markers in the mammary tissues. In situ detection of 8-OHdG, a commonly used oxidative stress marker, through immunohistochemistry (IHC) revealed that 8-OHdG levels in the mammary epithelial cells of *Aldh2-/-* mice were significantly increased (Fig. 4A), highlighting oxidative stress induced by ALDH2 deficiency. We next employed Western blotting to analyze key markers of cellular stresses and found that protein levels of phosphorylated p38 MAPK and NF-κB were significantly upregulated in the mammary tissues of *Aldh2-/-* mice compared to controls (Fig. 4B). Importantly, these changes were accompanied by the upregulation of Nrf2, a master regulator of antioxidant response [39]. This underscores the role of p38 MAPK, NF-κB, and Nrf2 activation in the interplay between oxidative stress signaling, inflammation, and antioxidant defense mechanisms in ALDH2 deficiency-induced cellular responses in mammary tissues. To further explore the mechanism of NF-κB activation, we examined whether ALDH2 knockout modulates expression of receptor activator of nuclear factor κB ligand (RANKL), a key regulator of osteoclast differentiation that has also been increasingly implicated in mammary stem cell regulation and breast oncogenesis [40]. Immunoblotting revealed that both the predominant full-length RANKL band and an additional lower–molecular weight band were elevated in *Aldh2-/-* mammary tissues, suggesting that ALDH2 deficiency enhances overall RANKL expression as well as the abundance of alternative isoforms or processed forms (Fig. 4B). To assess the association of the increased oxidative stress with DNA damage, we tested DNA damage signaling in the mammary tissues with ALDH2 KO. Protein levels of p53 and phosphorylated H2A.X, markers of DNA damage, were remarkably increased in *Aldh2-/-* tissues. Meanwhile, protein levels of oncogenic MDM2, a p53-specific E3 ubiquitin ligase responsive to p53 activation, were also significantly upregulated. These results demonstrate that ALDH2 deficiency-induced metabolic changes, even in the absence of alcohol exposure, are strong triggers for inducing oxidative stress that results in DNA damage in mammary tissues, which involves the activation of p38 MAPK, NF-κB and Nrf2 pathways.

**Figure 4.**
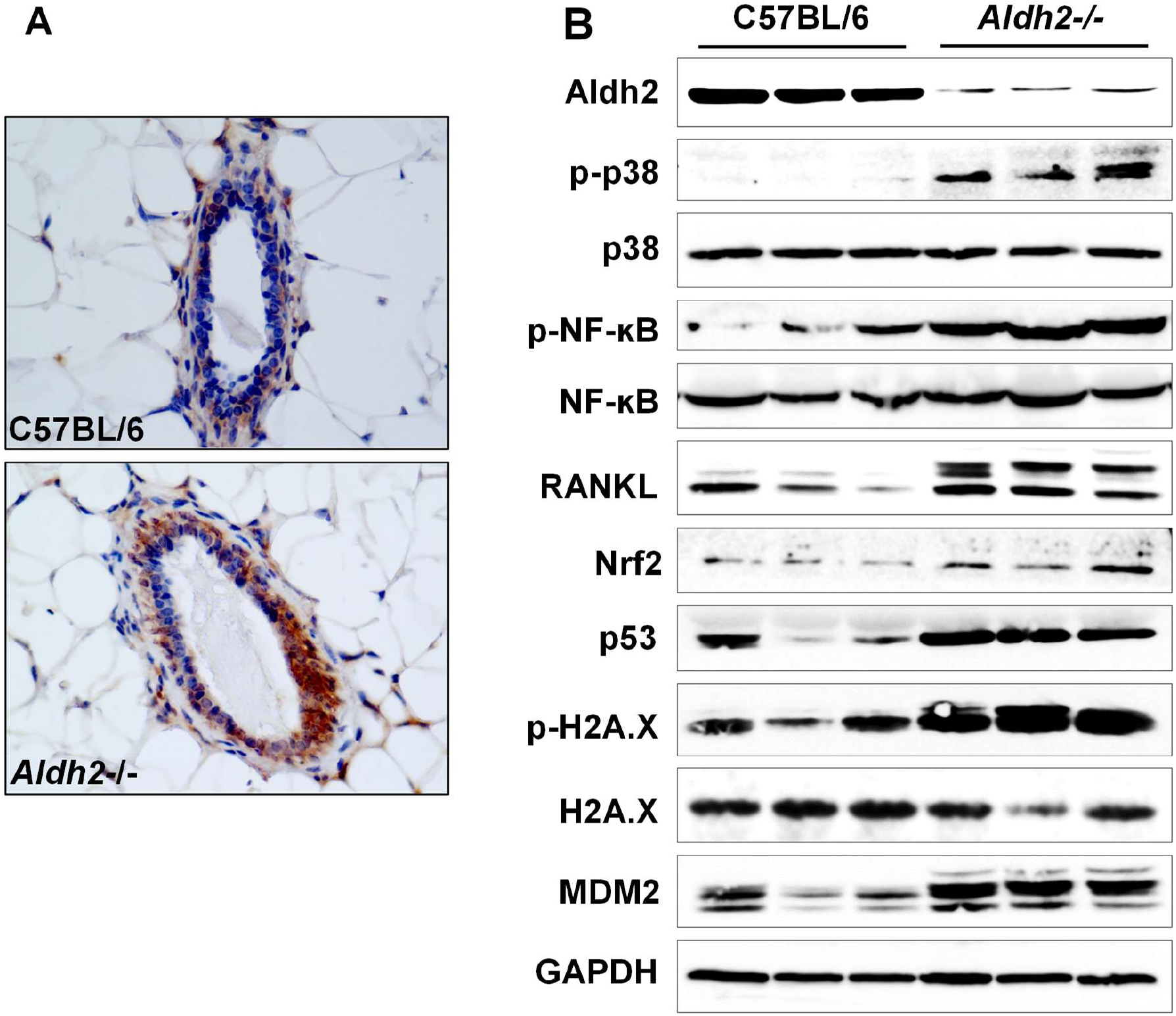
ALDH2 KO activates p38 MAPK and NF-κB-associated oxidative stress and induces DNA damage pathway in mammary tissues. **A)** Immunohistochemical staining of 8-OHdG (dark brown) in mammary tissues of control and *Aldh2-/-* mice. **B)** ALDH2 KO induces the expression and phosphorylation of key markers of oxidative stress and DNA damage pathways in mammary tissues of control and *Aldh2-/-* mice. Protein levels of indicated markers from three mice per group were detected with Western blotting.

### 3.5 ALDH2 KO induces hormonal regulation disruption and concomitant activation of erbB3-associated signaling in mammary tissues

Hormonal signaling is central to mammary development and oncogenesis [41]. We therefore examined the effect of ALDH2 KO on the signaling of the ER/PR pathway. Western blot analysis indicated that protein levels of ERα, PR, and phosphorylated ERα (Ser118) were significantly increased (Fig. 5A), suggesting that ALDH2 KO induces both the expression and phosphorylation/activation of ER signaling. Consistently, protein levels of ER targets, including c-Myc and Bcl-2, were also upregulated. These results indicate that ALDH2 deficiency strongly enhances ER signaling. To further support this novel finding, we examined *in situ* signals of phosphorylated ERα (S118) using IHC in the mammary tissues. The results showed that p-ERα (S118) levels were significantly increased in the *Aldh2-/-* tissues (Fig. 5B). Additionally, to further demonstrate ALDH2 KO-induced activation of ER signaling, we measured mRNA levels of several ER-target genes, including ESR1/ERα, PR, MYC, JUN, and BCL2. The results revealed that the mRNA levels of these ER-target genes in the mammary tissues of *Aldh2-/-* mice were significantly upregulated (Fig. 5C). Data from Western blot analysis, IHC staining, and qRT-PCR provide compelling *in vivo* evidence that ALDH2 deficiency induces ERα expression and activation in mammary tissues, potentially contributing to proliferative mammary development and epithelial cell repopulation.

**Figure 5.**
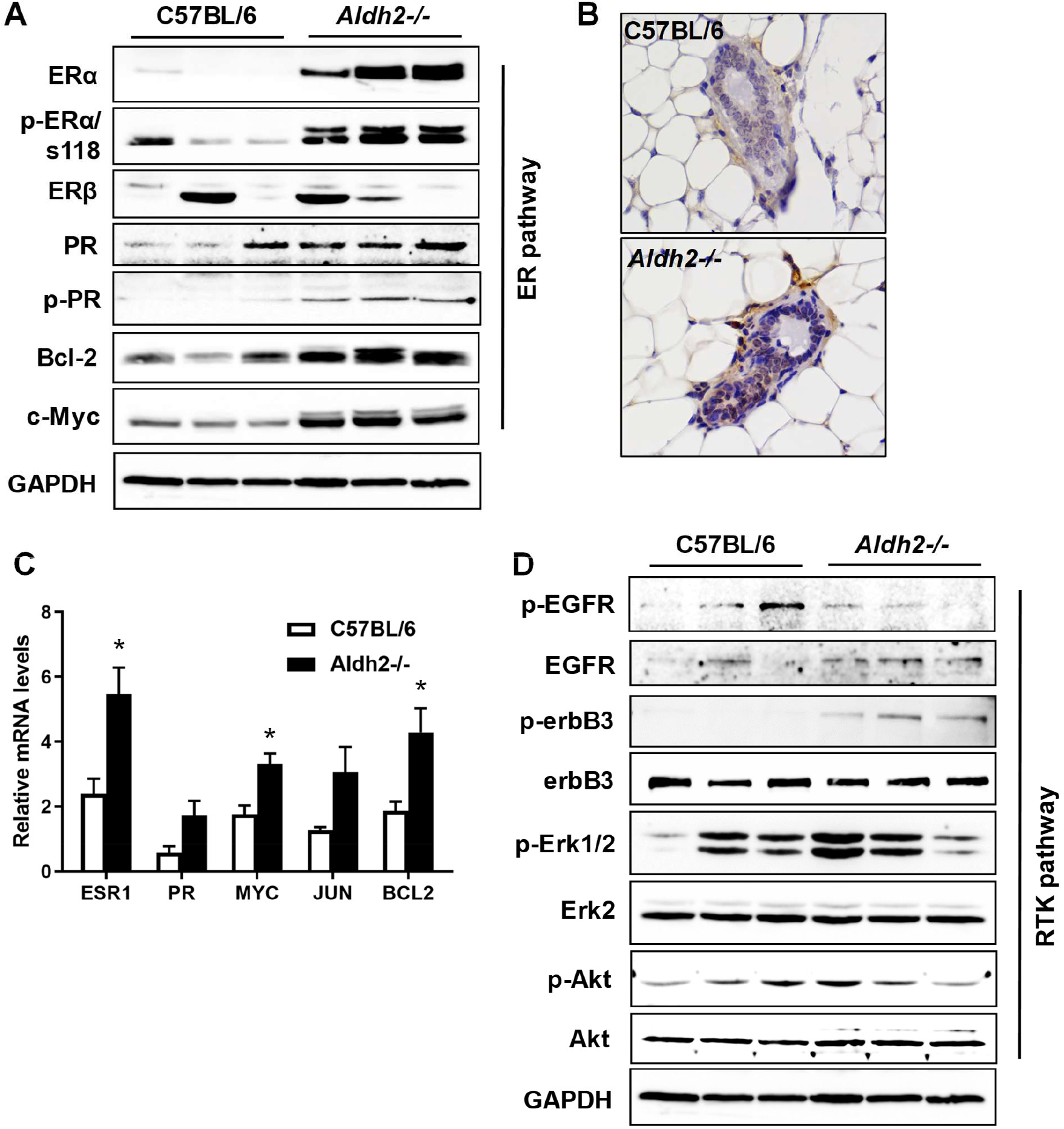
ALDH2 KO upregulates estrogen receptor signaling in conjunction with receptor tyrosine kinase pathways in mammary tissues. **A)** Western blot analysis of the expression and phosphorylation of ER and PR, along with selected target genes. Protein lysates from the mammary tissues of three mice in each group (control and *Aldh2-/-*) were separated using SDS-PAGE, followed by probing with specific antibodies against the indicated markers. **B)** IHC analysis of phosphorylated-ERα (S118) (brown color) in control and *Aldh2-/-* mammary tissues. **C)** Quantitative real-time PCR analysis of relative mRNA levels of selected ER target genes in mammary tissues of each group. Total RNA was extracted from mammary tissues, followed by RT-PCR detecting individual genes. The relative mRNA levels of individual genes were normalized with β-actin. (* p < 0.05). **D)** Western blot analysis of the expression and phosphorylation of key markers of EGFR associated signaling. Protein levels of indicated markers in control and *Aldh2-/-* mammary tissues were detected as described in (A).

Since ER signaling is closely regulated by its crosstalk with RTK signaling [42], we examined the protein levels of EGFR, erbB3, and downstream markers of Erk and Akt in the mammary tissues of both groups. Notably, the protein levels of erbB3 were distinctively upregulated, which was accompanied by increased levels of EGFR and phospho-Erk1/2 in the ALDH2 KO group (Fig. 5D). These data suggest that the activation of RTK signaling and the concomitant activation of ER signaling may interact to promote mammary epithelial cell proliferation and stemness.

### 3.6 ALDH2 KO induces striking upregulation of cyclin D1 and the activation of Rb-E2F1 pathway in mammary tissues

The Rb/E2F1-cyclin D1 axis drives cell cycle progression and proliferation [43]. To further characterize ALDH2 KO-induced proliferation of mammary epithelial cells, we examined the expression of cell cycle markers associated with proliferation, including Rb, E2F1, and cyclin D1. We found that ALDH2 KO significantly induced the phosphorylation of Rb protein and upregulated the protein levels of E2F1 (Fig. 6A). In particular, protein levels of cyclin D1 were remarkably increased. ALDH2 deficiency-induced upregulation of cyclin D1 was further supported by increased mRNA levels (Fig. 6B) and IHC staining of cyclin D1 in the mammary tissues (Fig. 6C). In light of these results, the data highlight cell cycle progression as a key target of ALDH2 KO-induced regulation. Since cyclin D1 is a target gene of ER signaling, the striking upregulation of cyclin D1 not only supports the activation of ER signaling in *Aldh2-/-* tissues but also suggests that cyclin D1 could serve as a marker for evaluating molecular changes induced by ALDH2 deficiency in mammary/breast tissues.

**Figure 6.**
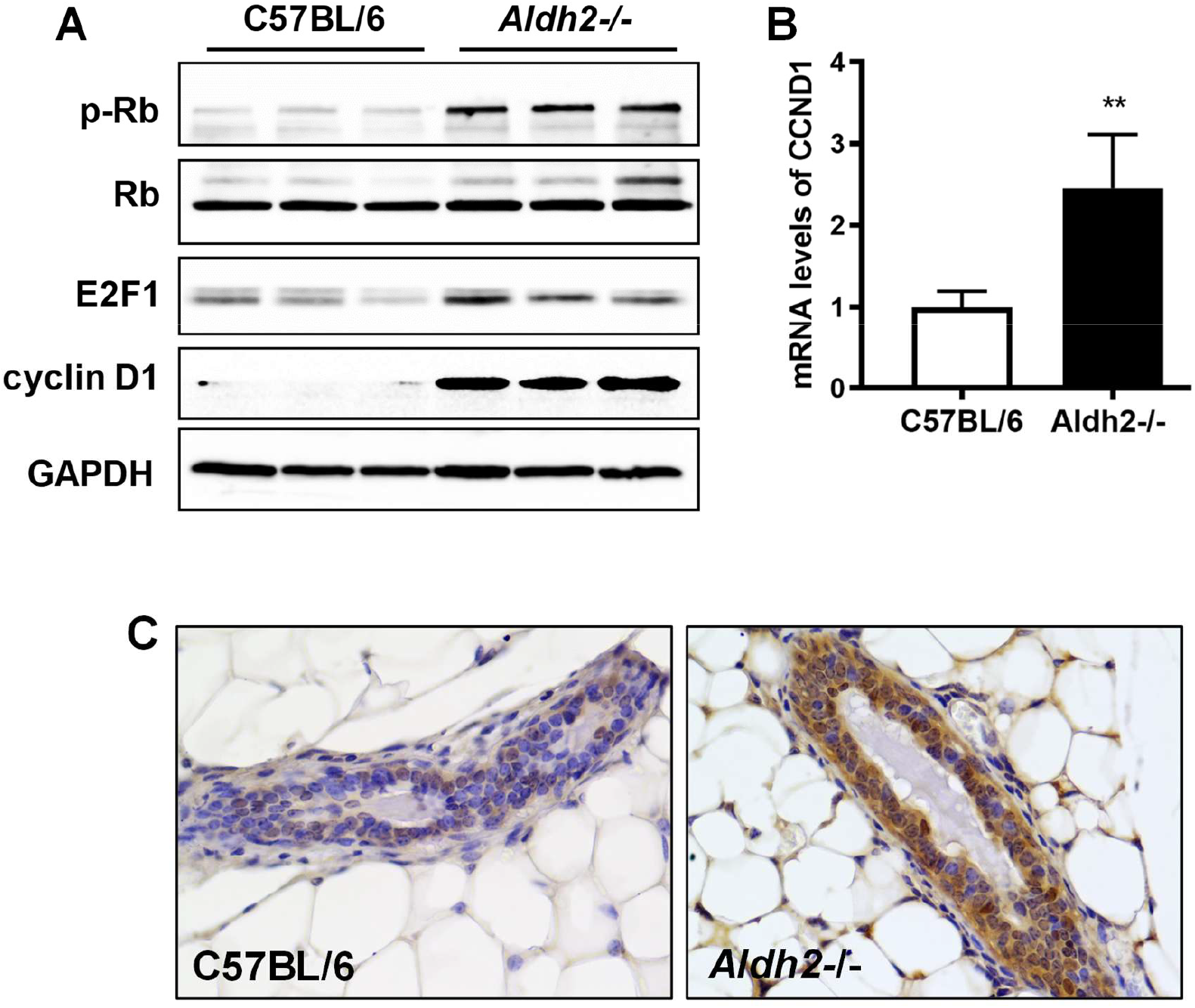
ALDH2 KO induces cyclin D1 upregulation and Rb-E2F1 pathway activation in mammary tissues. **A)** Protein levels of Rb, E2F1 and cyclin D1 in mammary tissues of control and *Aldh2-/-* mice. The indicated markers probed with specific antibodies were detected with Western blotting as in Fig. 4. **B)** Relative mRNA levels of CCND1 (cyclin D1) were quantified with real-time PCR as detailed with Fig. 5. **C)** Immunohistochemical analysis of cyclin D1 expression in mammary tissues of the paired groups. (** p < 0.01).

## 4. Discussion

The present study provides novel insights into the effects of ALDH2 deficiency on mammary gland development and its potential implications for breast cancer risk. As the first to report on studies of ALDH2 deficiency-associated mammary pathophysiology, our findings demonstrate that ALDH2 KO induces enhanced activation of oxidative stress and significant structural and functional changes in mammary tissues with molecular insight. In particular, the association of ALDH2 KO-induced ER signaling and proliferation with alterations in epithelial cell stemness could be pertinent to the understanding of breast cancer pathogenesis.

Our histopathological analysis indicates that ALDH2 KO leads to more complex ductal growth and increased ductal thickness in mammary glands compared to controls, as evidenced by whole mount and H&E staining (Fig. 1). This observation is supported by Ki67 staining, which revealed a higher percentage of proliferative cells in *Aldh2-/-* tissues. Moreover, our examination of the mammary whole mounts of 73-week old control and *Aldh2-/-* mice found that the mammary ductal trees of the *Aldh2-/-* mice showed a more complex ductal structure and epithelial density (unpublished data). The implication of enhanced mammary gland proliferation aligns with previous findings showing that alterations in metabolic enzymes can impact cell growth and tissue remodeling in other tissues [44]. The accumulation of endogenous toxic aldehydes or other metabolites due to ALDH2 deficiency may exacerbate oxidative stress within mammary tissues, contributing to the observed hyperplasia and increased cellular turnover.

Flow cytometry further revealed expansion of both luminal and basal epithelial cell populations in *Aldh2-/-* mice, based on CD24/CD49f analysis (Fig. 2). The basal subpopulation, which includes stem and progenitor cells crucial for mammary development and tumorigenesis [45], suggests increased stem cell activity and mammary epithelial cell proliferation. In parallel, the luminal subpopulations, composed of differentiated cells that line the inner layer of mammary ducts and alveoli [46], were also elevated, indicating enhanced differentiation or glandular activity. Additionally, we demonstrated that ALDH2 KO significantly increases progenitor cell-enriched P2 subpopulations in CD61/CD49f-based analysis (Fig. 2), which is signified by the essential role of luminal progenitor cells in both normal development and tumorigenesis [36, 37]. The observed epithelial cell repopulation and enrichment of potential progenitor cells suggest that ALDH2 deficiency influences not only the overall hierarchy of mammary epithelial cells but also the stemness of these cells, potentially expanding the pool of cells that could give rise to tumor initiation cells in the presence of other factors [32]. The above observed mammary epithelial cell repopulation is also supported by our assessment of mammary epithelial cell stemness in the *Aldh2-/-* model using colony-forming assays and mammosphere formation studies (Fig. 3). The increased colony numbers and enhanced mammosphere formation efficiency indicate that ALDH2 deficiency promotes stem cell characteristics among mammary epithelial cells. These novel findings underscore the potential contribution of ALDH2 deficiency-altered metabolic states and microenvironment to modified stem cell dynamics that may promote the potential emergence of tumor initiation cells. The alterations in the intrinsic regulation of mammary epithelial cells in this model system, along with the methodologies employed in this study, offer a solid basis and valuable resources for investigating ALDH2 deficiency-induced mammary tumorigenesis using other mammary tumor models.

The development and function of the mammary gland, as well as mammary epithelial cells, are sensitive to cellular stress and changes in the microenvironment [47]. Oxidative stress is one such factor that can significantly influence mammary gland development and the behavior of mammary epithelial cells [48, 49]. However, the consequences can vary depending on the specific induction factors and the intensity of the stress. It remains unknown whether ALDH2 deficiency-associated oxidative stress, in the absence of alcohol exposure, affects mammary development and epithelial cell regulation. Consistent with the observed phenotypic changes, our analysis of oxidative stress signaling and DNA damage in mammary tissues illuminates the intricate molecular mechanisms by which Aldh2 KO induces alterations in mammary epithelial stemness, particularly under oxidative stress conditions. The upregulation of the p38 MAPK and NF-κB pathways in the mammary tissues of ALDH2 KO mice signifies a robust activation of cellular stress responses (Fig. 4). The induction of Nrf2 underscores the activation of the antioxidant defense mechanism. Notably, while the involvement of oxidative stress markers such as p38 MAPK and NF-κB is well-documented in various cancer studies [50, 51], the specific impact of ALDH2 deficiency and the resulting endogenous toxic metabolites on mammary tissues and cells has not been thoroughly explored. Furthermore, our study uniquely demonstrates that the accumulation of these metabolites is sufficiently potent to induce significant DNA damage, as evidenced by increases in p53 and H2A.X levels, suggesting that ALDH2 deficiency not only signals cellular oxidative stress but also poses a risk for genomic instability. This is consistent with recent reports indicating that alcohol-induced or endogenous aldehydes impair adult tissue stem cells, such as hematopoietic, neural and intestinal stem cells [52-54].

Our study reveals for the first time that ALDH2 deficiency markedly upregulates RANKL in mammary tissues, linking aldehyde stress to RANKL-driven tumorigenesis [55]. While RANKL expression is known to be induced by hormonal cues such as progesterone and hypoxia [55, 56], its regulation by metabolic enzyme deficiency has not been reported. Notably, RANKL plays a critical role in mammary stem cell regulation [55, 57]. It also acts upstream of NF-κB and p38 MAPK signaling, thereby amplifying stress responses and stem-like features in breast cancer cells [58]. These findings place ALDH2 deficiency within this pathogenic network and suggest that toxic aldehyde accumulation may prime mammary epithelial cells for malignant transformation through RANKL induction and stress pathway activation.

ER signaling is crucial for normal mammary gland development and plays a significant role in breast cancer risk [59-61]. Our results indicate that ALDH2 deficiency significantly activates the ER signaling pathway in mammary tissues, as evidenced by upregulated levels of ERα and PR, along with enhanced phosphorylation of ERα (S118) and PR (S190) in mammary tissues with ALDH2 KO (Fig. 5). The upregulation of ER target genes, such as Bcl-2, cyclin D1, and c-Myc, along with IHC staining for p-ERα (S118), further supports the activation of ER signaling. The significant increase in cyclin D1 in ALDH2 KO tissues may contribute to enhanced signaling in the Rb/E2F1-cyclin D1 axis (Fig. 6). Given the pivotal role of ER and PR regulation in mammary epithelial proliferation and the maintenance of stemness [59], our results not only provide *in vivo* evidence of ALDH2 KO-mediated upregulation of ER signaling but also suggest a mechanistic role for ER/PR signaling dysregulation in ALDH2 deficiency-induced mammary epithelial cell proliferation and stemness. Additionally, we found that ALDH2 KO induced phosphorylation/activation of erbB3 and associated signaling (Fig. 5). ErbB3, a potent receptor tyrosine kinase (RTK) of the EGFR/erbB family, plays a critical role in mammary development and tumorigenesis [62]. The distinctive upregulation of phosphorylated erbB3 in *Aldh2-/-* tissues underscores its involvement in ALDH2-related changes in mammary epithelial cell proliferation and repopulation. Since ERα-S118 is a substrate of Erk1/2 [63], the enhanced activation of Erk1/2 and the increase in phosphorylated ERα-S118 highlight the connection between ER and RTK pathways in the alterations of mammary proliferation and development induced by ALDH2 deficiency. Notably, the activation of ER signaling and erbB3 due to ALDH2 deficiency has not been previously reported, which warrant further investigation. In the context of breast oncogenesis, our findings regarding the disruption of hormonal signaling associated with ALDH2 deficiency suggest an increased risk of breast cancer under these metabolic conditions.

Taken together, our research provides compelling evidence that ALDH2 deficiency promotes mammary epithelial cell proliferation and morphogenesis, which is associated with altered stem-like properties among mammary epithelial cells. By elucidating the role of ALDH2 in maintaining epithelial cell integrity and regulating stemness, we introduce a critical lens through which the developmental biology of the mammary gland can be understood in the context of breast cancer risk. Notably, these changes were observed in the *Aldh2-/-* model independent of alcohol exposure. Our data underscores the significance of the toxic impact of endogenous aldehyde accumulation or toxic metabolites on mammary development and potential risk for breast cancer in individuals with ALDH2 deficiency. Our study also highlights the interaction between metabolic dysregulation, such as oxidative stress, due to ALDH2 deficiency and its downstream effects on mammary tissue remodeling, cellular proliferation, and stemness. Our results from this model system lay a solid foundation for further investigation of ALDH2 deficiency-induced intrinsic metabolic and signaling pathways as vital contributors to breast cancer etiology. Critical questions have arisen from this study, such as the effects of ALDH2 KO on alcohol-induced mammary epithelial proliferation and stemness, as well as whether ALDH2 KO promotes tumorigenesis in mammary tumor models. These questions are being addressed in our ongoing projects.

In conclusion, our investigation highlights the multifaceted impact of ALDH2 deficiency on mammary gland development and its potential implications for breast cancer risk. The alterations in cell proliferation, coding for stemness, and activation of oxidative stress, hormonal modulation, and RANK/RANKL signaling pathways bolster the consideration of ALDH2 as a critical player in mammary biology. As the understanding of genetic and metabolic interactions in breast cancer becomes increasingly intricate, further research into the molecular underpinnings of ALDH2 deficiency and its association with cancer risk will be essential for developing targeted prevention strategies in susceptible populations. By elucidating the mechanisms through which ALDH2 influences mammary development, future studies may uncover therapeutic targets for the intervention in breast cancer.

## CRediT authorship contribution statement

**Zhikun Ma**: Data curation, formal analysis, writing – review & editing; **Amanda B Parris**: Data curation, formal analysis, writing - review & editing; **Miles Lester**: Data curation; **De’ja Gissendanner**: Data curation; **Vasilis Vasiliou**: transgenic animal model and manuscript critics; **Xiaohe Yang**: Conceptualization, funding acquisition, investigation, project administration, writing – original draft

